# DECR1 is an androgen-repressed survival factor that regulates PUFA oxidation to protect prostate tumor cells from ferroptosis

**DOI:** 10.1101/865626

**Authors:** Zeyad D. Nassar, Chui Yan Mah, Jonas Dehairs, Ingrid J.G. Burvenich, Swati Irani, Margaret M. Centenera, Raj K. Shrestha, Max Moldovan, Anthony S. Don, Andrew M. Scott, Lisa G. Horvath, David J. Lynn, Luke A. Selth, Andrew J. Hoy, Johannes V. Swinnen, Lisa M. Butler

**Affiliations:** University of Adelaide Medical School and Freemasons Foundation Centre for Men’s Health, University of Adelaide, Adelaide, SA 5005, Australia; South Australian Health and Medical Research Institute, Adelaide, SA 5000, Australia; KU Leuven-University of Leuven, LKI-Leuven Cancer Institute, Department of Oncology, Laboratory of Lipid Metabolism and Cancer, Leuven, B-3000, Belgium; Tumour Targeting Laboratory, Olivia Newton-John Cancer Research Institute, and School of Cancer Medicine, La Trobe University, Melbourne, VIC 3084, Australia; Dame Roma Mitchell Cancer Research Laboratories, University of Adelaide, Adelaide, SA 5005, Australia; NHMRC Clinical Trials Centre, and Centenary Institute, The University of Sydney, Camperdown, NSW 2006, Australia; Garvan Institute of Medical Research, Darlinghurst, NSW 2010; University of Sydney, Camperdown, NSW 2006; and University of New South Wales, Sydney, NSW 2052, Australia; College of Medicine and Public Health, Flinders University, Bedford Park, SA 5042, Australia; Discipline of Physiology, School of Medical Sciences, Charles Perkins Centre, Faculty of Medicine and Health, The University of Sydney, Camperdown, NSW 2006, Australia

**Keywords:** Fatty acid oxidation, PUFA, DECR1, Patient-derived explants, Prostate cancer

## Abstract

Fatty acid β-oxidation (FAO) is the main bioenergetic pathway in prostate cancer (PCa) and a promising novel therapeutic vulnerability. Here we demonstrate therapeutic efficacy of targeting FAO in clinical prostate tumors cultured *ex vivo,* and identify *DECR1,* which encodes the rate-limiting enzyme for oxidation of polyunsaturated fatty acids (PUFAs), as robustly overexpressed in PCa tissues and associated with shorter relapse-free survival. *DECR1* is a negatively-regulated androgen receptor (AR) target gene and, therefore, may promote PCa cell survival and resistance to AR targeting therapeutics. DECR1 knockdown in PCa cells selectively inhibited β-oxidation of PUFAs, inhibited proliferation and migration of PCa cells, including treatment resistant lines, and suppressed tumor cell proliferation *in vivo*. Mechanistically, targeting of DECR1 caused cellular accumulation of linoleic acid, enhanced mitochondrial oxidative stress and lipid peroxidation, and ferroptosis. These findings implicate PUFA oxidation via DECR1 as a previously unexplored facet of FAO that promotes survival of PCa cells.

## Introduction

Prostate cancer (PCa) is the most prevalent male cancer and the second leading cause of cancer deaths in men in Western societies (1). For patients with locally-recurrent and/or metastatic disease, androgen deprivation therapy (ADT) has remained the frontline strategy for clinical management since the 1940s (2), due to the dependence of PCa cells on androgens for growth and survival. Although ADT is initially effective in most patients, ultimately all will relapse with castration resistant prostate cancer (CRPC), which remains incurable. The failure of ADT is attributed to the emergence of adaptive survival pathways that reprogram androgen signaling and/or activate alternative tumor survival pathways. Consequently, the development and FDA approval of agents that more effectively target androgen signaling, including enzalutamide (ENZ, Xtandi; an AR antagonist) (3–5), has expanded the therapeutic options for CRPC. Nevertheless, even these approaches cannot durably control tumor growth and there is considerable variability in the nature and duration of responses between different patients (3,6,7). Thus, alternative therapeutic strategies that enhance response to ADT, and thereby prevent or delay PCa progression to CRPC, are essential.

Increasingly, targeting cancer cell metabolism is a focus of research efforts (8). While fundamental differences in cellular metabolism pathways between normal and malignant cells were detected by Warburg in the 1920s (9), clinical targeting of cancer metabolism has not kept pace with the research advances in understanding metabolic features of cancer cells. PCa is mainly dependent on lipid metabolism for energy production (10). The overexpression of genes involved in lipid metabolism is characteristic of PCa at both early and advanced stages (11–17), while recent proteomic analyses of primary PCa and bone metastases have shown clear associations between levels of lipid metabolic enzymes, PCa initiation and progression (18, 19). These observations suggest that PCa may be particularly amenable to metabolic targeting strategies. Despite this, the role and complexity of lipid/fatty acid (FA) metabolism in PCa and its potential as a target for therapy remains underexplored, particularly in the context of a more complex tumor microenvironment.

Until recently, most attention has focused on the therapeutic targeting of *de novo* FA synthesis and, most recently, uptake of FAs in PCa to limit their availability as a source of energy and cell membrane phospholipids (14,15,20). However, it has become evident in work from our group and others that β-oxidation of FAs, as the ultimate fate of FAs in the energy production cycle, is upregulated in PCa cells, stimulated by a lipid-rich extracellular environment and critical for viability (10,21,22). In this study, we evaluated the targeting of FA β-oxidation (FAO) in patient-derived prostate tumor explants (PDE) to provide the first clinically-relevant evidence that targeting this pathway is efficacious. We subsequently identify DECR1, a rate-limiting enzyme in an auxiliary pathway for polyunsaturated fatty acid (PUFA) β-oxidation, as a promising novel therapeutic vulnerability for PCa. Importantly, we show that DECR1 is an androgen-repressed gene induced in PCa cells in response to ADT and/or AR-targeted therapies, implicating PUFA oxidation as an adaptive survival response that may contribute to emergence of CRPC and treatment resistance.

## Results

### Targeting FA oxidation is efficacious in patient-derived PCa explants

In addition to our recent report of enhanced FAO in PCa cells (22), an accumulating body of evidence supports the efficacy of targeting key enzymes involved in FAO using *in vitro* and *in vivo* models of PCa (23–25). However, to date there is limited evidence that targeting this pathway would be clinically efficacious, which prompted us to target this pathway in clinical tumors. Using our well-defined patient derived explant (PDE) model that recapitulates the complexity of the clinical tissue microenvironment (26), we targeted the rate-limiting enzyme in mitochondrial FAO, carnitine palmitoyltransferase-1 (CPT-1), in cultured PDEs using the chemical inhibitor etomoxir. Consistent with literature reports, etomoxir had weak activity against the LNCaP PCa cell line *in vitro*, with an IC50 of 170 µM (Figure S1A), but was considerably potent in the PDEs, in which a dose of 100 µM inhibited cell proliferation by an average of 48.4% ± 16.6% (n=13 patients; *p*<0.05) (***Figure 1A***). Etomoxir effectively inhibited FAO in the tissues, evidenced by a significant decrease in multiple acylcarnitines in the conditioned medium (***Figure 1B***).

**Figure.1.**
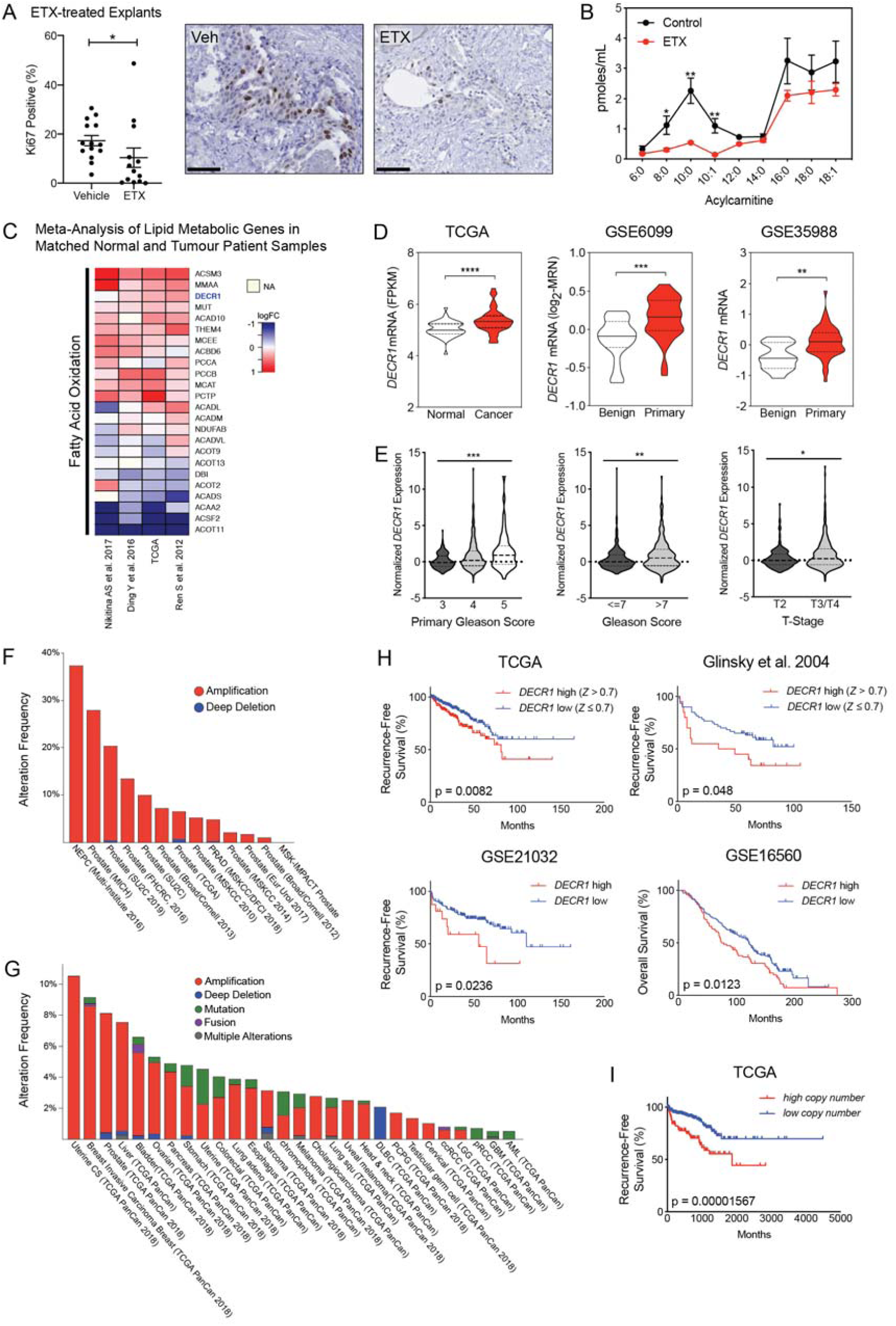
Fatty acid β-oxidation genes are overexpressed in prostate cancer and targeting this process is effective in patient-derived human prostatic *ex vivo* tumor explants. **(A)** Etomoxir reduced cell proliferation in patient-derived human prostatic *ex vivo* tumor explants. Tissues were treated with 100 µM etomoxir for 72 hours, sections were fixed in formalin, paraffin embedded and stained against the proliferative marker Ki67 (n=13) (scale bar = 50 µm). **(B)** Etomoxir (100 µM) decreased acylcarnitine species, the products of CPT1 activity. Acylcarnitines secreted to the conditioned medium were measured after 72 hours treatment of PDEs (n=9). (**C)** A meta-analysis of fatty acid oxidation genes using four clinical datasets with malignant and matched normal RNA-sequencing data (n=122). Genes were rank-ordered on the basis of their *meta effect size* scores in PCa malignant tissues versus matched normal tissues. **(D)** Violin plots demonstrate DECR1 mRNA overexpression in PCa primary/malignant tissues compared to normal/benign tissues in three independent datasets. **(E)** DECR1 mRNA expression is associated with PCa primary gleason score, total gleason score (>7) and diseases stage (T-stage). Data were extracted from TCGA PCa dataset. **(F)** Histogram displaying DECR1 mutation and copy-number amplification frequency across 13 PCa genomic datasets, and **(G)** across 28 tumor types. Histograms were obtained from CbioPortal platform. **(H)** DECR1 mRNA expression is associated with shorter relapse-free survival in TCGA PCa, Glinsky et al. (30) and GSE21032 datasets, and shorter overall survival rates in GSE16560 dataset. **(I)** DECR1 copy number amplification frequency is associated with shorter relapse-free survival in TCGA PCa dataset. Data in **(A)** are represented as as the mean ± s.e.m and were statistically analysed using a Wilcoxon matched-pairs signed rank test. Data in **(B)** are represented as the mean ± s.e.m and were statistically analysed using two-tailed Student’s *t*-test. Data in **(D)** and **(E)** are represented as violin plots in GraphPad prism: the horizontal line within the violin represents the median, and were statistically analysed using a Mann-Whitney two-tailed t-test. Data in **(H)** and **(I)** were statistically analysed using a two-sided log-rank test. \**p*<0.05, \*\**p* <0.01, \*\*\**p* <0.001 and \*\*\*\**p*<0.0001.

In order to prioritize key functional genes involved in PCa progression, we conducted a meta-analysis of the expression of 735 genes involved in lipid metabolism (as identified from REACTOME) in four clinical datasets with malignant and matched normal RNA sequencing data (27–29). Genes were rank-ordered on the basis of their *meta effect size* scores in PCa malignant versus matched normal tissues (Figure S1B). The meta-analysis revealed a strikingly consistent deregulation of lipid metabolism genes, including genes involved in FAO (***Figure 1C***), despite the predicted high inter-individual heterogeneity of patient PCa tissues. We conducted disease-relapse survival analysis using TCGA data for each of the top 20 genes from the meta-analysis. This identified *DECR1* as a robustly overexpressed gene in PCa tissues that is associated with shorter relapse-free survival rates.

### DECR1 is upregulated in clinical prostate tumors

Consistently, *DECR1* mRNA expression was significantly higher in malignant compared to benign prostate tissues in ten independent expression datasets of PCa tissues versus non-malignant tissues (***Figure 1D***, S1C). Further analysis of the TCGA data revealed increased *DECR1* expression with increased Gleason score or in with advanced disease stage (***Figure 1E***). Consistent with our observation of increased DECR1 mRNA expression in PCa, *DECR1* gene copy gain was evident in several clinical datasets accessed via cBioPortal (31) (***Figure 1F***). Interestingly, the top three cancer types exhibiting increased *DECR1* copy gain were hormone-dependent tumors (uterine, breast and prostate), suggestive of a relationship between DECR1 expression and hormone signaling (***Figure 1G***). *DECR1* mRNA expression was associated with shorter relapse-free survival rates and overall survival rates (***Figure 1H***), and in the TCGA dataset, *DECR1* amplification was significantly associated with shorter recurrence-free survival rates (***Figure 1I***).

We confirmed overexpression of DECR1 protein in clinical PCa using two independent proteomic datasets (***Figure 2A***). We observed overexpression of DECR1 in PCa tissues (n=8) compared with benign tissues (n=3) (***Figure 2B***), and increased expression was evident in high grade versus low grade cancer tissue. Quantitative IHC staining analysis revealed a significant increase of DECR1 expression in malignant *vs* benign tissues. Furthermore, intra-tissue analysis exposed a significant increase of DECR1 expression in malignant regions *vs* benign ones within the same core (***Figure 2C***). DECR1 expression was markedly increased in a panel of hormone-dependent and -independent cancer cell lines compared with non-malignant PNT1 and PNT2 prostate cell lines (***Figure 2D***). Consistent with its function, DECR1 localises to the mitochondria, confirmed using immunocytochemistry and Western blot of nuclear, cytoplasmic and mitochondrial cell fractions (***Figure 2E***). Together, the mRNA and protein findings suggest that expression of DECR1 is closely linked to PCa progression and patient outcomes, and therefore might represent an unexplored therapeutic target.

**Figure.2.**
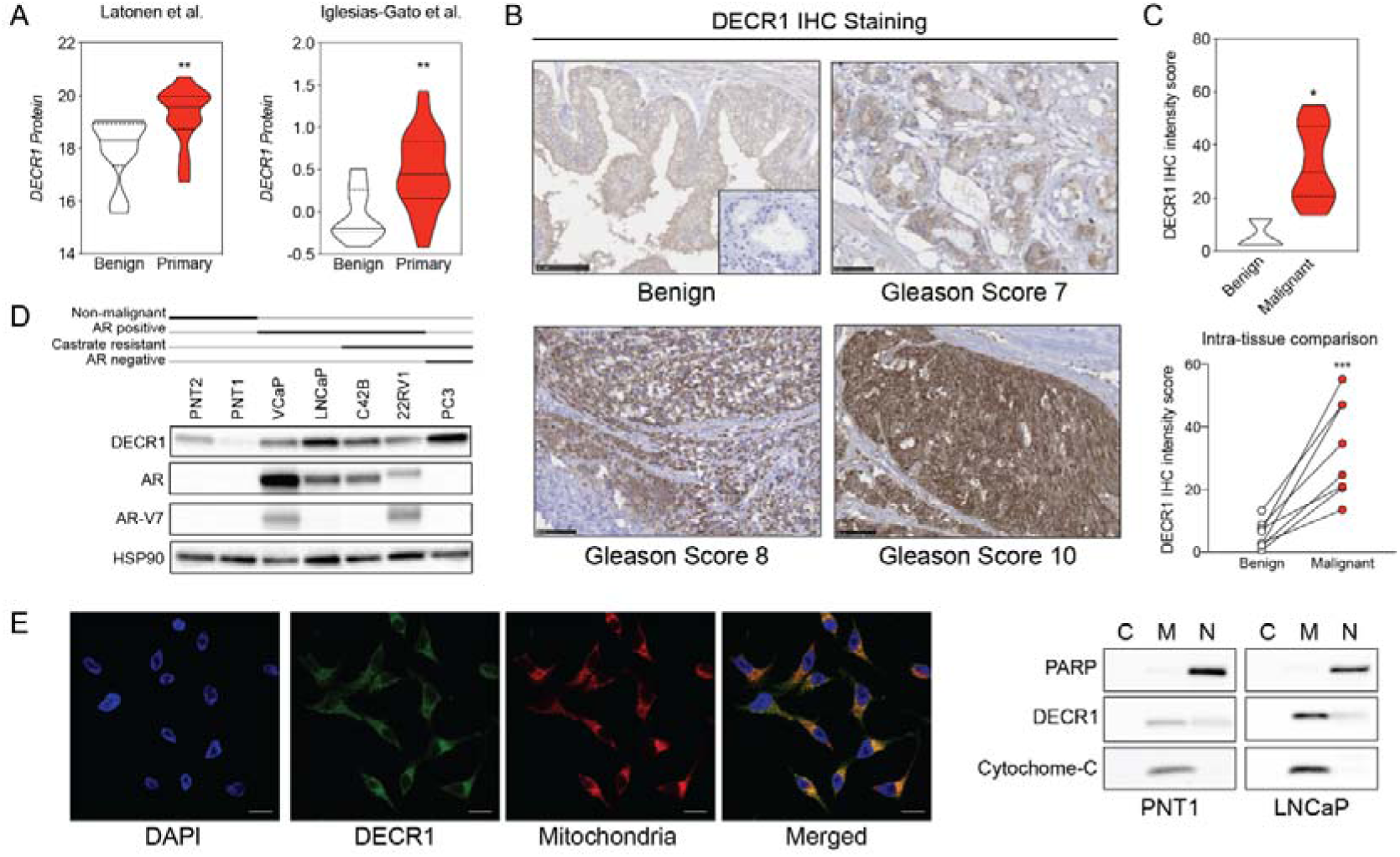
DECR1 protein in overexpressed in prostate malignant cells/tissues. **(A)** Violin plots of DECR1 protein overexpression in primary PCa tissues compared to benign prostate tissues in two independent datasets. **(B)** Representative DECR1 IHC staining of benign prostate tissues and PCa tissues (negative control stain included in bottom right box). Scale bar, 50 µm. **(C)** Violin plot of DECR1 protein expression in a validation cohort consisting of benign prostate tissues (n=3) and PCa tissues (n=8) (*top panel*). Intra-tissue IHC analysis of DECR1 expression in PCa tissues (n=8) (*bottom panel*). **(D)** DECR1 protein expression in non-malignant prostate cell lines (PNT1 and PNT2) and PCa cell lines (LNCaP, VCaP, 22RV1, C42B and PC3). **(E)** Immunocytochemistry staining of LNCaP cells to determine the subcellular localization of DECR1: nuclei were labelled using DAPI; mitochondria were labelled using MitoTracker Red; and DECR1 proteins were labelled using Alexa Fluor 488 secondary antibody, (Scale bar =10 μm). Immunoblot of PNT1 and LNCaP cells separated into cytosolic, mitochondrial and nuclear fractions and incubated with poly (ADP-ribose) polymerase (PARP) and cytochrome-C antibodies to mark nuclear and mitochondrial fractions. Data are represented as violin plots in GraphPad prism: the horizontal line within the violin represents the median. Statistical analysis was performed using a Mann-Whitney two-tailed t-test (**A** and **C** *top panel*), or two-tailed paired t-test (**C** *bottom panel*): \**p*<0.05, \*\**p* <0.01 and \*\*\**p* <0.001.

### DECR1 is a directly androgen-repressed gene in PCa

The relationship between androgen receptor (AR) signaling and lipid metabolic genes is well established. Many studies have reported a marked stimulatory effect of AR on key lipid metabolism pathways either directly or indirectly through activation of a family of transcription factors called sterol regulatory element-binding proteins (SREBPs) (32). We therefore investigated the relationship between AR and DECR1 using a panel of *in vitro*, *ex vivo* and *in vivo* models. DECR1 expression is notably more abundant in AR-negative cells (PC3) than in AR-expressing cells (***Figure 2D***), consistent with negative regulation of DECR1 expression by AR. We confirmed that androgen (dihydrotestosterone) significantly decreased DECR1 expression in androgen-dependent LNCaP and VCaP cell lines at both mRNA and protein levels (***Figure 3A***). Data mining of publicly available microarray datasets also revealed downregulation of DECR1 in LNCaP cells after treatment with DHT or the synthetic androgen R1881 (Figure S2A; GSE7868, GSE22606). In contrast to the effect of androgens, AR-targeted therapies increase DECR1 expression. LNCaP and VCaP cell treatment with the androgen antagonist enzalutamide (ENZ) significantly increased DECR1 expression at both mRNA and protein levels (***Figure 3B***). *In vivo*, LNCaP tumors exhibit increased DECR1 expression in mice treated with ENZ (10 mg/kg) or castration, which was enhanced further in mice treated with both ENZ and castration (***Figure 3C***). Our observations were supported by published microarray datasets which showed that treatment of LNCaP or VCaP cells with ENZ increases *DECR1* mRNA expression (Figure S2B; GSE69249). *In vivo*, castration of mice increased DECR1 expression in prostate (Figure S2C; GSE5901), while in LNCaP/AR xenografts treatment with the AR antagonist ARN-509 (apalutamide) for 4 days significantly increased DECR1 expression (Figure S2D; GSE52169). To confirm androgenic regulation of DECR1 in a clinical context, we validated these data using PDEs. ENZ treatment of PDEs significantly increased DECR1 expression whereas, as expected, mRNA levels of the well-characterized AR target genes *KLK3* and *KLK2* were decreased (***Figure 3D, E***). To determine whether AR directly represses *DECR1*, we interrogated published chromatin immunoprecipitation (ChIP) sequencing data. In VCaP cells, AR bound strongly to the *DECR1* promoter in response to DHT treatment, but not when co-treated with AR antagonists (***Figure 3F***; GSE55064). Moreover, AR binding was enriched at the *DECR1* promoter in benign and malignant prostate tissues (Figure S2E; GSE56288). A site-specific ChIP-qPCR assay revealed DHT-stimulated AR occupancy at this region in LNCaP cells (***Figure 3G***). Collectively, these data reveal *DECR1* as a novel AR-repressed gene.

**Figure.3.**
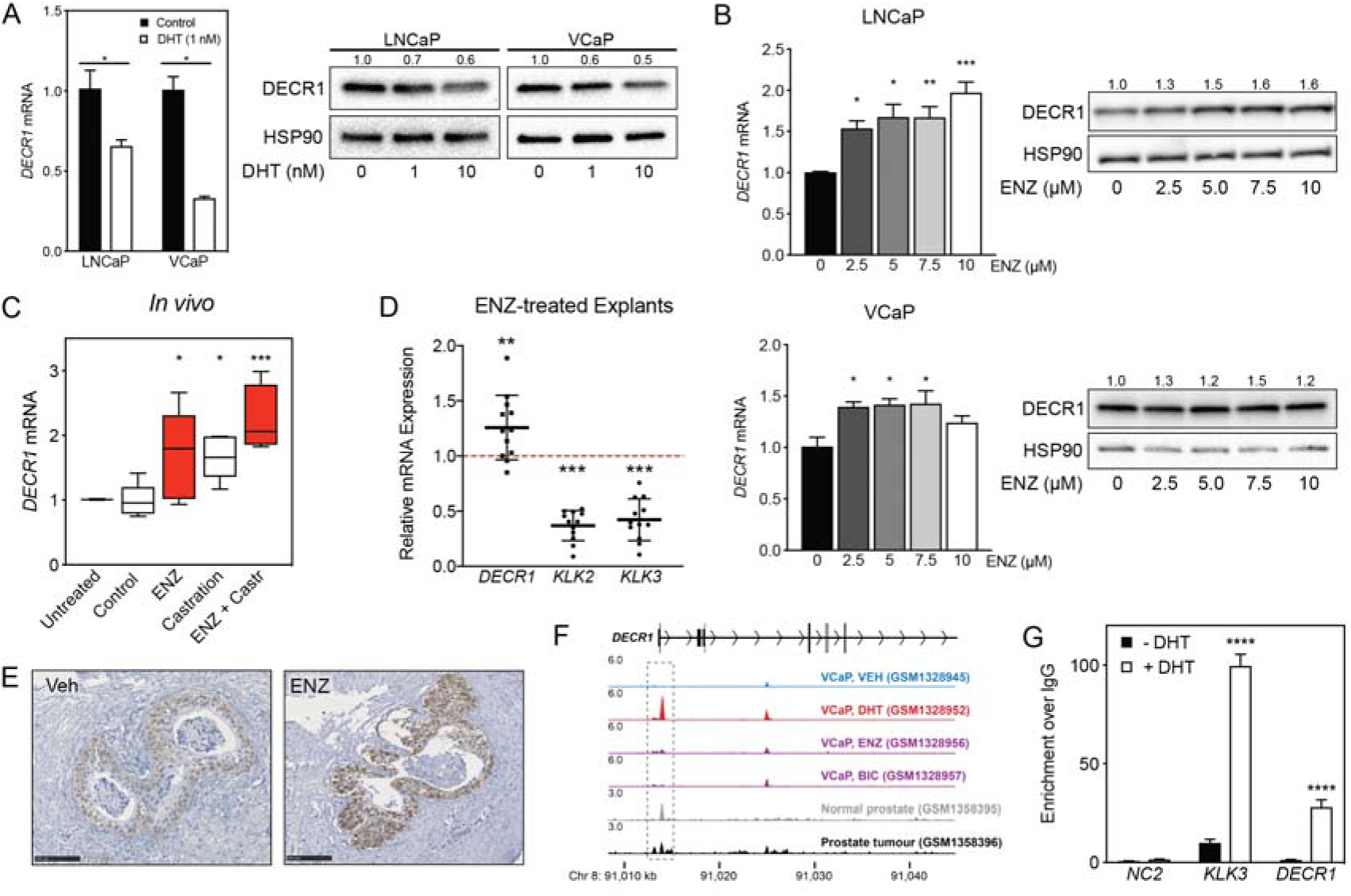
DECR1 is androgen-repressed gene. **(A)** DECR1 mRNA and protein was measured by qRT-PCR and western blot analysis after PCa cell treatment with dihydrotestosterone (DHT), or **(B)** enzalutamide (ENZ). Relative mRNA expression of *DECR1* was calculated using comparative CT method, where the cells treated with vehicle (control) were set to 1 and normalised to the geometric mean CT value of *GUSB* and *L19* (housekeeping genes). Densitometry quantification of relative DECR1 protein expression was normalized to the HSP90 internal control. *n* = 3 independent experiments. **(C)** qRT-PCR analysis of *DECR1* mRNA expression in LNCaP-derived tumors treated with enzalutamide (ENZ) and/or castration (Castr). **(D)** qRT-PCR analysis of *DECR1* and androgen regulated genes *KLK2* and *KLK3* mRNA expression in a cohort of 10 patient-derived human prostatic *ex vivo* tumor explants treated with enzalutamide (ENZ, 10 µM). Relative mRNA expression was calculated using comparative CT method, where the matched untreated tissue from the same patient was set to 1 and normalized to the geometric mean CT value of *TUBA1B, PPIA* and *GAPDH*. **(E)** Representative DECR1 IHC staining of patient-derived human prostatic *ex vivo* tumor explants treated with enzalutamide (ENZ, 10 µM). Scale bar, 100 µm. **(F)** AR ChIP-sequencing data from VCaP cells (top panel), normal human prostate and primary human prostate tumor specimens (bottom panel). Data from GSE55064 and GSE56288. **(G)** ChIP-qPCR analysis demonstrates AR binding at *DECR1* locus in LNCaP cells after treatment with DHT. Data in bar graphs (**A, B** and **G)** are represented as the mean ± s.e.m. Data in **(C)** are represented as box plots using the Tukey method in GraphPad prism. Statistical analysis was performed using two-tailed Student’s *t*-test (**A, C, D** and **G**) or one-way ANOVA, followed by Dunnett’s multiple comparisons test **(B)**: \**p*<0.05, \*\**p* <0.01, \*\*\**p* <0.001 and \*\*\*\**p*<0.0001.

### Targeting DECR1 disrupts PUFA oxidation in PCa cells

FAs are metabolized mainly in mitochondria through the β-oxidation process to generate acetyl-CoA, which enters the tricarboxylic acid cycle (TCA) and produces ATP and NADH as energy for the cell. Unlike saturated FAs, all unsaturated FAs with double bonds originating at even-numbered positions, and some unsaturated FAs with double bonds originating at odd-numbered positions, require three auxiliary enzymes to generate intermediates that are harmonious with the standard β-oxidation pathway (33, 34): Enoyl CoA isomerase (ECI1), 2,4 Dienoyl-CoA reductase (DECR1) and Dienoyl CoA isomerase (ECH1) (***Figure 4A***). DECR1 catalyses the rate limiting step in this pathway (35). Given the critical role of DECR1 in PUFA metabolism, we studied the consequences of DECR1 downregulation on β-oxidation of PUFAs in PCa cells. DECR1 knockdown was achieved successfully (>80% downregulation) using 2 different siRNAs (***Figure 4B***). DECR1 knockdown resulted in an increase in linoleic acid (***Figure 4C***) as well as an accumulation of 2-trans,4-cis-decadienoylcarnitine (acylcarnitine 10:2; ***Figure 4D***), an intermediate of linoleic acid metabolism, indicating incomplete PUFA β-oxidation.

**Figure.4.**
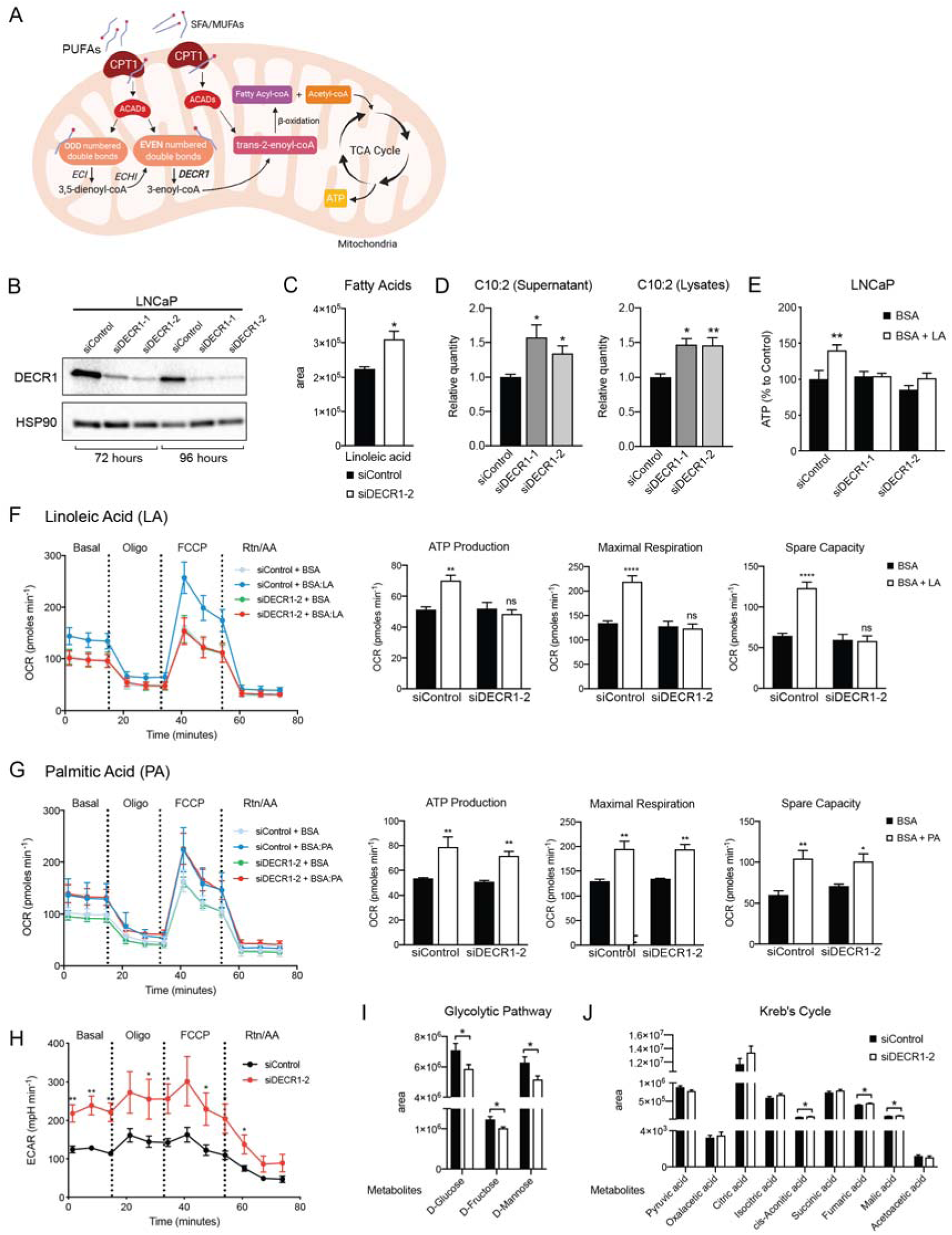
DECR1 knockdown interrupts PUFA β-oxidation in PCa cells. **(A)** Schematic of DECR1 function in fatty acid (FA) β-oxidation. In order to translocate FAs into the mitochondria, CPT1 converts long-chain acyl-CoA species to their corresponding long-chain acylcarnitine species. This is followed by a dehydrogenation step mediated by acyl CoA dehydrogenase (ACAD) to generate trans-2-enoyl-CoA, the only intermediate that can be processed by downstream enzymes in the β-oxidation process. Many FAs have unsaturated bonds either on an odd-numbered carbon or in the cis-configuration, resulting in the generation of enoyl-CoA intermediates that cannot be directly processed via the downstream β-oxidation enzymes. These FAs require the activity of 3 auxiliary enzymes, ECI1, ECH1 and DECR1 in order to form trans-2-enoyl-CoA before undergoing β-oxidation. DECR1 catalyzes the conversion of either 2-trans,4-cis-dienoyl or 2-trans,4-trans-dienoyl-CoA to 3-trans-enoyl-CoA. A complete cycle of β-oxidation results in the release of the first two carbon units as acetyl-CoA, and a fatty-acyl-CoA minus two carbons. The acetyl-CoA enters the TCA cycle to produce energy (ATP). The shortened fatty-acyl-CoA is processed again starting with the ACADs to form trans-2-enoyl-CoA either directly or with the aid of the auxiliary enzymes depending on the presence of double bonds. This process continues until all carbons in the fatty acid chain are turned into acetyl-CoA. **(B)** DECR1 protein expression after 72 hours or 96 hours siRNA transfection. Densitometry quantification of relative DECR1 protein expression was normalized to the HSP90 internal control. **(C)** Linoleic acid level in LNCaP cells quantified in following 96 hours DECR1 knockdown using GC QQQ targeted metabolomics. **(D)** Relative quantities of the C10:2 acylcarnitine species in LNCaP cell conditioned medium (left) or cell lysates (right) (n=3). **(E)** Quantification of ATP levels in LNCaP cell lysates. LNCaP cells were transfected with DECR1 siRNAs for 48 hours and then starved in no-glucose medium and treated with the lipolysis inhibitor DEUP (100µM) in the presence (BSA-LA) or absence (BSA) of the PUFA linoleic acid for 48 hours before measuring ATP levels. (**F)** Oxygen consumption rate (OCR) was assessed in LNCaP cells supplemented with the PUFA linoleic acid (LA) or **(G)** the saturated fatty acid palmitic acid (PA). Each data point represents an OCR measurement. ATP production, maximal mitochondrial respiration and mitochondrial spare capacity were assessed. **(H)** Extracellular acidification rate (ECAR) was assessed in LNCaP cells. Each data point represents an ECAR measurement. For experiments **(F-H)** LNCaP cells were transfected with DECR1 siRNAs for 72 hours, then starved in substrate limited medium for 24 hours; the assay was run in FAO assay medium. **(I & J)** Metabolites were quantified in LNCaP cells following 96 hours DECR1 knockdown using GC QQQ targeted metabolomics. Data in bar graphs are represented as the mean ± s.e.m (n=3). Statistical analysis was performed using two-tailed Student’s *t*-test: \**p*<0.05, \*\**p* <0.01 and \*\*\*\**p* <0.0001.

Mitochondrial β-oxidation provides reducing equivalents that drive ATP production. LNCaP cells increased ATP levels in response to exogenous linoleic acid supplementation when cultured in glucose-free media containing the lipase inhibitor diethylumbelliferyl phosphate (DEUP), to prevent the cells from using intracellular FAs (***Figure 4E***). However, cells transfected with DECR1-targeting siRNAs failed to increase ATP levels with linoleic acid supplementation (***Figure 4E***), indicating impaired capacity to metabolize PUFAs.

Next, we employed extracellular flux analysis to determine the intrinsic rate and capacity of PCa cells to oxidise PUFAs in conditions where other exogenous substrates were limiting. Exogenous linoleic acid stimulated basal oxygen consumption rates (OCR), as a measure of mitochondrial oxidative phosphorylation, and maximal respiration, ATP production, and mitochondrial spare capacity (***Figure 4F***, S3A) as determined by consecutive cell exposure to respiration chain inhibitors and uncouplers. This supports the observed increased in total ATP levels in response to linoleic acid supplementation (***Figure 4E***). Importantly, DECR1 knockdown prevented the exogenous linoleic acid induction of basal and maximal respiration, ATP production, and mitochondrial spare capacity (***Figure 4F***, S3A). In contrast, DECR1 knockdown has no impact on mitochondrial metabolism of the saturated FA, palmitate (***Figure 4G***). Further, DECR1 knockdown increased glycolysis, as determined by ECAR (***Figure 4H***, S3B), as well as decreased glucose and fructose concentrations (***Figure 4I***), to sustain TCA cycle intermediate levels (***Figure 4J***) as a consequence of disruption of PUFA β-oxidation. Collectively, these results demonstrate the DECR1 is critical for PUFA metabolism in LNCaP PCa cells.

### Targeting DECR1 suppresses PCa oncogenesis

The consistently increased expression of DECR1 in PCa tissue and its association with shorter-relapse times and survival rates (***Figure 1 & 2***), taken together with its impact on PUFA metabolism (***Figure 4***), suggested that it may contribute to PCa cell viability and invasive behaviour. We evaluated the impact of DECR1 downregulation or overexpression on various oncogenic properties of PCa cells using a panel of *in vitro* and *in vivo* experiments. While there was no effect of DECR1 downregulation on the non-malignant prostate cell line PNT1, a significant attenuation of PCa proliferation and induction of cell death was observed in a panel of PCa lines (***Figure 5A***), comprising androgen-dependent (VCaP and LNCaP), CRPC (22RV1 and V16D) and acquired ENZ-resistant cells (MR49F). Notably, this effect on PCa cell viability was lost when cells were cultured in lipid-depleted media (Figure S4), suggesting that the observed effect is due to interference with FA metabolism. In contrast, stable DECR1 overexpression significantly enhanced LNCaP cell viability (***Figure 5B***) and colony formation ability (***Figure 5C***), while stable DECR1 knockdown using a short hairpin vector markedly decreased colony formation (***Figure 5D***). DECR1 knockdown also decreased LNCaP growth in 3D spheroids (***Figure 5E***), which better mimic *in vivo* conditions than 2-dimensional cell culture (36). In addition, DECR1 knockdown reduced LNCaP, 22RV1 and MR49F cell migration by ∼ 50% (***Figure 5F***) and 22RV1 invasion by ∼65% (***Figure 5G***). LNCaP cells stably depleted of DECR1 showed highly variable growth rates *in vivo*, but inspection of the resultant tumors revealed significantly reduced cellular proliferation compared to control cells, concomitant with reduced DECR1 expression (***Figure 5H, I***).

**Figure.5.**
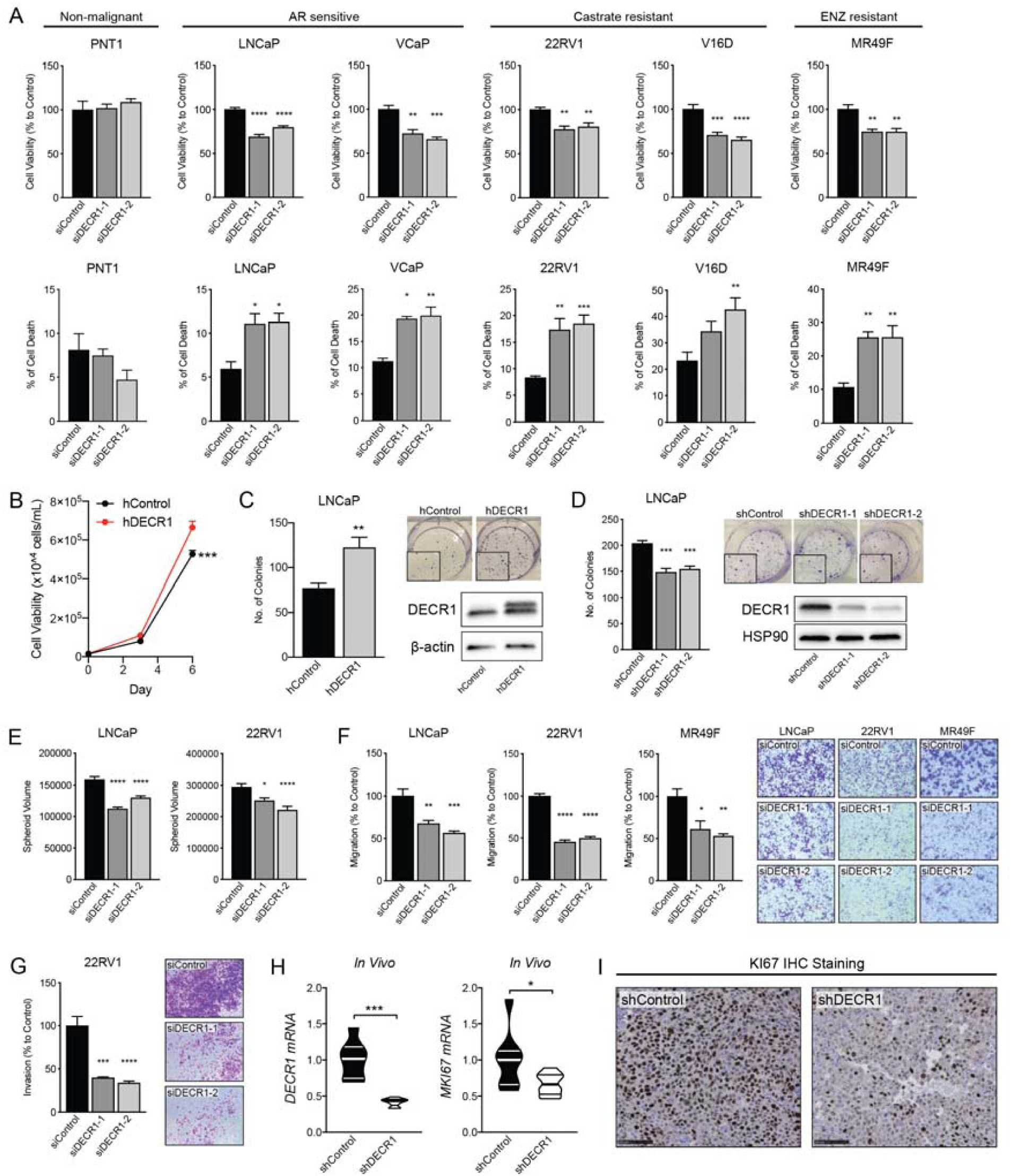
DECR1 knockdown suppresses PCa cell growth. **(A)** Cell viability after DECR1 knockdown in non-malignant PNT1 prostate cells; hormone-responsive PCa cell lines (LNCaP and VCaP); castrate-resistant V16D and 22RV1 cell lines and enzalutamide-resistant MR94F cells cultured in full serum media. **(B)** Cell viability of stable DECR1-overexpressed LNCaP cells cultured in full serum media. Cell viability and cell death were measured using trypan blue exclusion following 96 hours DECR1 knockdown. Percentages are represented relative to the control siRNA; *n* = 3 independent experiments per cell line. **(C)** Clonogenic cell survival of LNCaP cells were assessed using colony formation assay. Stable DECR1-overexpressed cells or **(D)** stable DECR1 knockdown was achieved using two different short hairpin (sh) vectors and DECR1 expression was confirmed using western blot. Cells were cultured for 2 weeks, washed with PBS, fixed with paraformaldehyde and stained with 1% crystal violet for 30 minutes. Colonies with more than 50 cells were counted manually; data shown is representative of *n* = 2 independent experiments. **(E)** LNCaP and 22RV1 cell growth in 3D spheres. Spheroids were prepared using the hang drop assay following 48 hours DECR1 knockdown. Spheroid volumes were determined after five days of culturing the cells in 20 µl drops; at least 25 spheres per cell line were assessed using the ReViSP software, *n* = 3 independent experiments per cell line. **(F)** LNCaP, 22RV1 and MR49F cell migration and **(G)** 22RV1 cell invasion were assessed using transwell migration/invasion assay. Cells were transfected with DECR1 siRNA or control siRNA for 48 hours. Equal number of cells were transferred to the upper inserts in serum free medium; lower chambers were filled with medium containing 5% serum as a chemoattractant. Plates were incubated for a further 48 hours. Migrated/invaded cells on the lower face of the inserts were washed with PBS, fixed with paraformaldehyde, stained with 1% crystal violet for 30 minutes, and counted manually; data shown is representative of *n* = 3 independent experiments. **(H)** Violin plots of *mKi67* and *DECR1* mRNA expression in LNCaP tumors (n = 5 mice, shControl; n = 4 mice, shDECR1). **(I)** Representative KI67 IHC staining of LNCaP tumors. Scale bar, 100µm. Data in bar graphs are represented as the mean ± s.e.m. Statistical analysis was performed using one-way ANOVA, followed by Dunnett’s multiple comparisons test: \**p*<0.05, \*\**p* <0.01, \*\*\**p* <0.001 and \*\*\*\**p*<0.0001.

### DECR1 targeting induces lipid peroxidation and cellular ferroptosis

DECR1 knockdown resulted in inhibition of PUFA β-oxidation and led to accumulation of PUFA (***Figure 4***), which are highly susceptible to peroxidation, so we next assessed the effect of DECR1 knockdown on lipid peroxidation. DECR1 knockdown increased levels of malondialdehyde, a marker of lipid peroxidation (37) (***Figure 6A***). In contrast, DECR1 overexpression markedly decreased cellular malondialdehyde levels (***Figure 6A***). We observed enhanced mitochondrial oxidative stress measured using MitoSOX, a mitochondrial superoxide indicator (***Figure 6B***), in response to DECR1 knockdown, while DECR1 overexpression significantly decreased mitochondrial oxidative stress under basal (***Figure 6C***) and linoleic acid-induced conditions (***Figure 6D***). Lipid peroxidation is a major driver of ferroptosis, an iron-dependent non-apoptotic form of cell death (38). Cell treatment with the ferroptosis inhibitor, ferrostatin, abolished the effect of DECR1 knockdown on PCa cell death (***Figure 6E***), while cell treatment with the ferroptosis inducer, erastin, enhanced the cytotoxic action of DECR1 downregulation (***Figure 6F***). Collectively, these results suggest that DECR1 expression protect cells from oxidative stress and that DECR1 knockdown-induced cell death is mediated, at least partly, by induction of ferroptosis.

**Figure.6.**
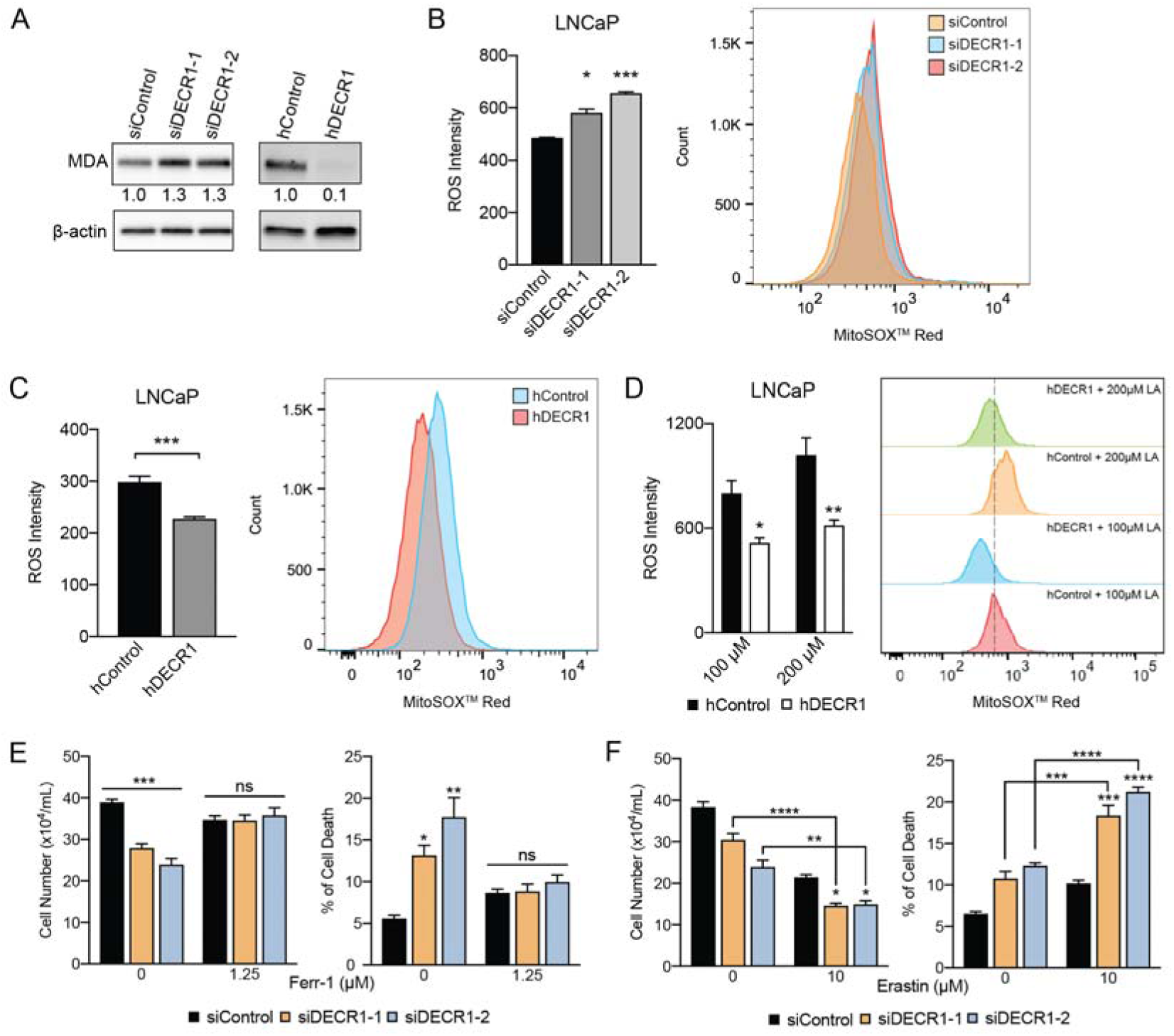
DECR1 knockdown induces PUFA accumulation, lipid peroxidation and ferroptosis. **(A)** MDA level, an oxidative stress marker, was measured by western blot in LNCaP cells transfected with DECR1 siRNAs. **(B)** Mitochondrial superoxide levels were quantified following 96 hours DECR1 knockdown using MitoSOX™ red stain. Fluorescent intensity was quantified using flow cytometry and ROS levels were presented as mean fluorescent intensity. **(C)** Mitochondrial superoxide levels were quantified in stable DECR1-overexpressed LNCaP cells using MitoSOX™ red stain under basal or **(D)** linoleic acid (100 µM or 200 µM) treated cells. Fluorescent intensity was quantified using flow cytometry and ROS levels were presented as mean fluorescent intensity. **(E)** Cell viability of LNCaP cells after 48 hours DECR1 knockdown, treated with ferrostatin (Ferr-1, 1.25 µM), or **(F)** erastin (10 µM). Data in bar graphs are represented as the mean ± s.e.m. Statistical analysis was performed using two-tailed Student’s *t*-test **(B-D)** or one-way ANOVA, followed by Holm-Sidak’s multiple comparisons test **(E and F)**: ns, not significant, \**p*<0.05, \*\**p* <0.01, \*\*\**p* <0.001 and \*\*\*\**p*<0.0001.

## Discussion

Metabolic rewiring is both a hallmark feature of cancer cells and a promising therapeutic vulnerability. Several anabolic and catabolic metabolism pathways have been explored, however, few agents have been investigated clinically. This can at least partly be explained by the relatively recent appreciation of cancer metabolism as a target, the high toxicity, particularly hepatotoxicity, expected to be associated with targeting certain metabolic pathways, the predictable metabolic heterogeneity within and between patients and the lack of intermediate pre-clinical models that can predict clinically efficacious outcomes. Previous research has largely focused on studying and targeting FA synthetic pathways in PCa. Major lipogenic enzymes such as ATP citrate lyase (ACLY), acetyl-CoA carboxylase (ACC) and fatty acid synthase (FASN) are all overexpressed in PCa compared to benign tissue (11,39,40). While many first-generation FA synthesis inhibiters (e.g. FASN inhibitors) showed promising preclinical efficacy against PCa, unfavourable drug solubility and pharmacokinetics profiles, off-target effects and side effects including weight loss have hindered clinical development of this agent class (14). In addition to *de novo* synthesis of FAs, PCa cells depend on lipid uptake from the circulation, and from stromal adipocytes (20,41,42). We showed previously that extracellular FAs are the major contributor to lipid synthesis in PCa (22). Moreover, targeting FA uptake using an antibody against CD36, a major transporter for exogenous FAs into the cells, reduced cancer severity in patient-derived xenografts, and *CD36* deletion slowed cancer progression in prostate-specific *PTEN^-/-^* mice. However, it is increasingly evident that PCa exhibits plasticity in attaining FAs and that crosstalk between *de novo* synthesis and FA uptake requires dual targeting of the two pathways to achieve maximal efficacy (20), an approach that would likely be associated with greater toxicity. In this study, we focused on another understudied aspect of FA metabolism in PCa, β-oxidation, to evaluate its therapeutic potential.

We showed previously (22), as have others (24, 43), that PCa cells exhibit increased FAO compared to prostate epithelial PNT-1 cells, or benign epithelial cells BPH-1 and WPMY-1. This metabolic phenotype is a vulnerability for PCa cells (21,23,25). FAO inhibitors such as etomoxir, perhexiline or ranolazin inhibited tumor growth *in vitro* and *in vivo* (21,23,25) and sensitized cells to ENZ treatment (23, 25). However, all of these studies were undertaken using immortalized cell line models of PCa, and as cancer metabolism is markedly influenced by the tumor microenvironment (44–46), employing preclinical models and primary tissues that retain the complexity of this microenvironment as a stepping stone to clinical trials may accelerate clinical translation and avoid futile targeting strategies or agents. In this study, we evaluated the efficacy of the FAO inhibitor, etomoxir, using our established PDE model. Remarkably, etomoxir inhibited effectively cell proliferation in PDEs (***Figure 1A***), strengthening the case that targeting FA oxidation may be a promising clinical strategy. Interestingly, etomoxir was more potent in inhibition of cell proliferation in PDEs than *in vitro* 2-dimensional growth of LNCaP cells, emphasising that *in vitro* models based on 2D cultured cells alone are sub-optimal when evaluating anti-metabolism agents. This problem is compounded by the fact that standard growth media is rich in sugar and proteins, but contains low levels of lipids. As the clinical development of etomoxir was terminated due to severe hepatotoxicity associated with treatment (47), we sought to identify new β-oxidation targets in PCa. As DECR1 is an androgen-repressed gene, its expression increases after castration or treatment with anti-androgens and is hypothesized to maintain tumor cell survival under castration conditions. Androgen-repressed genes are markedly understudied compared with androgen-induced genes, despite the fact that they possibly mediate adaptive survival pathways when androgen signalling is perturbed, and have already yielded novel therapeutic targets (48, 49).

Surprisingly, the biological role of DECR1 in PCa has never been examined, and very little is known about its function in other cancers. Human DECR1 deficiency is lethal, with patients exhibiting hypocarnitinemia, decreased cellular oxygen consumption, increased lactic acidosis, and unusual accumulation of FA intermediates in urine and blood due to incomplete β-oxidation (50, 51). DECR1-null mice exhibit impaired lipid metabolism, hypoglycaemia and activation of ketogenesis, and cold intolerance (52, 53). These phenotypes highlight the critical role of DECR1 in lipid metabolism. We confirmed the metabolic activities of DECR1 in PCa cells using a panel of metabolomic and lipidomic analyses. DECR1 knockdown increased levels of certain acylcarnitine species, indicating inhibition of β-oxidation. These results are consistent with previous studies reporting acylcarnitine accumulation in *Decr1^-/-^* mice and *DECR1*-deficient patients (50, 52). We showed that DECR1 knockdown selectively inhibited PUFA catabolism, accompanied by an increase in glycolysis rate, which is also consistent with previous reports of FAO inhibition or impaired mitochondrial function leading to enhanced glucose uptake and glycolysis (43, 51). Lipidomic analysis showed that DECR1 knockdown increased the abundance of PUFAs and certain lipids, particularly PE and PI phospholipid species, accompanied by increased levels of mitochondrial oxidative stress and particularly lipid peroxidation. In contrast to MUFAs, PUFAs are highly susceptible to peroxidation, thereby enhancing free radical generation and accumulation of toxic lipid peroxides (54, 55). Consistent with a role for PUFA oxidation in DECR1 function, ectopic DECR1 overexpression decreased mitochondrial oxidative stress. An important cellular protective response to excess intracellular lipid peroxides is the induction of ferroptosis, an iron-dependent form of cell death that is triggered by lipid peroxidation. Here, we show that the ferroptosis inhibitor ferrostatin or the ferroptosis inducer erastin abolished and augmented the effect of DECR1 on PCa cell death, respectively. These findings support a mechanism by which DECR1 knockdown-induced cell death is a ferroptosis-mediated process caused by PUFA accumulation. It is therefore possible that PCa cells commonly select for DECR1 overexpression, not only to enhance ATP production to fulfil energy requirements, but also to protect cells from the tumoricidal effects of excess PUFAs.

PUFAs are essential FAs that cannot be synthesized in mammals and are only obtained from the diet. Dietary fat not only promotes obesity, but also PCa progression and disease aggressiveness. Several preclinical PCa studies have compared high-fat (or Western-style diets) versus low fat diet and reported that the former promotes AKT and ERK activity, tumor growth, tumor incidence in genetically engineered/transgenic mouse models, tumor progression to CRPC and metastasis (reviewed in (56)). Clinical case-control studies indicated saturated fat is associated with an increased risk of advanced PCa (57–60). Several underlying mechanisms were proposed to explain the association between dietary fat and PCa development and progression, including growth factor signalling (such as IGF-1), inflammation, and endocrine modulation (56). While the evidence supporting the negative impacts of saturated dietary FAs are more consistent, the effect of dietary PUFAs on PCa aggressiveness remains inconclusive and differences between omega-3 and omega-6 have been reported (57,58,61,62). In contrast to omega-3 PUFAs, which are reported to inhibit PCa progression (63), omega-6 PUFAs (64, 65), and higher omega-6/omega-3 PUFA ratio, increase PCa risk (66, 67). Dietary intervention by decreasing total fat intake and increasing omega-3 PUFAs was found to improve PCa survivorship (68–71). It is unclear whether there is a preference for n-6 or n-3 PUFA β-oxidation in PCa cells, however both require DECR1 for complete β-oxidation.

Although FAO is a complex process that requires the activity of several enzymes, to date the entire focus of drug development strategies has been inhibitors against CPT1, the rate-limiting step of FA β-oxidation. CPT1 is responsible for synthesising fatty acyl-carnitines from fatty acyl-CoAs which are then transported from the cytoplasm into the mitochondria by carnitine acylcarnitine translocase for subsequent processing then entry into the β-oxidation pathway. Even though targeting CPT1 is efficient in inhibiting FA β-oxidation, the clinical use of CPT1 inhibitors is challenging. Based on our current findings, we propose that DECR1 is an attractive alternative target to CPT1. CPT1 inhibition would suppress β-oxidation of all long FA species (saturated FA, MUFA and PUFA), whilst in contrast DECR1 is specific for PUFA. Homozygous *CPT1* deficiency, of either the liver or muscle isoform, is lethal in mice (72–74), but *Decr1^−/−^* mice are viable, and clinical symptoms arose only after metabolic stress (52, 53). The marked overexpression of DECR1 in prostate tumors across multiple clinical cohorts, potentially coupled to PUFA-related dietary interventions, may lend further selectivity to targeting strategies. Of note, the crystal structure of DECR1 active site has been solved (35), and thus developing DECR1 inhibitors is feasible.

In summary, herein we strengthen the evidence base for the critical importance of FAO and, specifically, PUFA oxidation in PCa, thereby identifying a promising new therapeutic candidate, DECR1.

## Materials and Methods

### Meta-Analysis of Lipid Metabolism Genes

Individual RNA-sequencing (RNA-seq) datasets composed of matched normal versus tumor prostate cancer patient tissue samples were acquired and are listed as follows: (1) The Cancer Genome Atlas (TCGA, n=53); (2) Nikitina AS et al. (GSE89223, n=10) (27); (3) Ren S et al. (n=14) (28); and (4) Ding Y et al. (GSE89194, n=45) (29). Before the meta-analysis, RNA-seq data was quality controlled and analysed using the R limma voom-*eBayes pipeline* (75).Effect sizes (log-fold changes) and corresponding variances were collected from the differential expression analysis under the matched-pairs design. Meta-analysis was performed by applying a restricted maximum-likelihood estimator (REML) within a random-effects model using the *rma* function from the R *metafor* package. At most one missing observation out of four was allowed per gene. Next, the retained gens were intersected with the list of pre-selected 735 genes involved in lipid metabolism as identified from REACTOME. Finally, the remaining genes were rank-ordered on the basis of their *meta effect size* scores across all four RNA-seq datasets. Top 20 candidate genes were selected for further disease-free survival association analyses from well-characterized clinical cohorts.

### Clinical Datasets

Transcriptomic data was downloaded from The Cancer Genome Atlas (TCGA) data portal, cBioPortal (31), and GEO; GSE21032 (76); GSE35988 (77); GSE6099 (78); GSE16560 (79). Proteomics data was extracted from Iglesias-Gato et al. (19).

### Cell lines and tissue culture

The human normal prostate epithelial cell lines PNT1 and PNT2, and prostate carcinoma cells LNCaP, VCaP, and 22RV1 were obtained from the American Type Culture Collection (Rockville, MD, USA). Castrate resistant V16D and enzalutamide resistant MR49F cells were derived through serial xenograft passage of LNCaP cells (80) were a kind gift from Prof. Amina Zoubeidi laboratory. Cell lines were verified in 2018 via short tandem repeat profiling (Cell Bank Australia). Cell lines were maintained in RPMI-1640 medium containing 10% fetal bovine serum (FBS; Sigma-Aldrich, NSW, Australia) in 5% CO_2_ in a humidified atmosphere at 37°C. Prior to androgen treatment, cells were seeded in medium supplemented with 5% dextran charcoal coated FBS (DCC-FBS) and after 24 hours, 1nM or 10nM of dihydrotestosterone (DHT) was added. For anti-androgen treatment, cells were cultured in growth medium supplemented with 2.5 µM, 5 µM, 7.5 µM or 10 µM Enzalutamide (dissolved in dimethyl sulfoxide, DMSO; Sigma Aldrich). The sources and experimental conditions for primary antibodies used in this study are listed in Supplementary Table 1. Primers were obtained from Sigma-Aldrich and their sequences are detailed in Supplementary Table 2 and 3.

### *Ex Vivo* culture of human prostate tumors

Patient derived-explant culture was carried out according to techniques established in our laboratory and as described previously (81–83). 6 mm/8 mm biopsy cores were collected from men undergoing robotic radical prostatectomy at St. Andrew’s Hospital (Adelaide, South Australia) with written informed consent through the Australian Prostate Cancer BioResource. The tissue was dissected into smaller 1 mm^3^ pieces and cultured on Gelfoam sponges (80 × 125 mm Pfizer 1205147) in 24-well plates pre-soaked in 500 µl RPMI 1640 with 10% FBS, antibiotic/antimycotic solution. Etomoxir (100 µM) or Enzalutamide (10 µM) was added into each well and the tissues were cultured in 5% CO_2_ in a humidified atmosphere at 37°C for 48 hours, then snap frozen in liquid nitrogen and stored at −80°C, or formalin-fixed and paraffin-embedded.

### Immunohistochemistry (IHC)

Paraffin-embedded tissue sections (2-4µm) were deparaffinized in xylene, rehydrated through graded ethanol, and blocked for endogenous peroxidase before being subjected to heat-induced epitope retrieval (81). IHC staining was performed using DECR1 (HPA023238, Sigma Aldrich, diluted 1:500) antibody and the 3,3′-Diaminobenzidine (DAB) Enhanced Liquid Substrate System tetrahydrochloride (Sigma Aldrich) as described previously (81). Intensity of DECR1 immunostaining was measured by video image analysis (81).

### Western Blotting

Protein lysates were collected in RIPA lysis buffer (10 mM Tris, 150mM NaCl, 1mM EDTA, 1% Triton X-100, 10% protease inhibitor). Western blotting on whole cell protein lysates were performed as previously described (81). *Cell Fractionation.* Protein lysates from each subcellular fraction (cytoplasm, mitochondria, and nucleus) were obtained from PNT1 and LNCaP cells using the cell fractionation kit (Abcam, VIC, Australia) according to the manufacturer’s protocol.

### Quantitative Real-Time PCR (qPCR)

RNA was extracted from cells using the RNeasy RNA extraction kit (Qiagen), followed by the iScript™ cDNA Synthesis kit (Bio-Rad, NSW, Australia). qPCR was performed with a 1:10 dilution of cDNA using SYBR Green (Bio-Rad) on a CFX384 Real-Time System (Bio-Rad). Relative gene expression was calculated using the comparative Ct method and normalized to the internal control genes *GUSB* and *L19* for prostate cancer cells and LNCaP-derived tumors, or *TUBA1B, PPIA* and *GAPDH* for PDEs.

### Analysis of published ChIP-seq data

AR ChIP-seq data from published external datasets, GSE56288 (clinical specimens; 7 normal prostate and 13 primary tumors) (84) and GSE55064 (VCaP cell line; Veh, DHT treated, MDV3100 treated and Bicalutamide treated) (85) were obtained from GEO and visualized using the Integrated Genome Browser (IGV).

### Chromatin Immunoprecipitation (ChIP)

LNCaP cells were seeded at 3 × 10^6^ cells/plate in 15 cm plates in RPMI-1640 medium containing 10% DCC-FBS for 3 days, then treated for 4 hours with 10 nM DHT or Vehicle (ethanol). AR ChIP was performed as described previously (86).

### Transient RNA interference

The human DECR1 ON-TARGET plus small interfering RNAs (siRNAs) and control siRNA (D-001810-01-20 ON-TARGET plus Non-targeting siRNA #1) were purchased from Millennium Science (VIC, Australia). Four siRNA were tested and the two most effective were selected for our experimentation: siDECR1-1 (J-009642-05-0002) and siDECR1-2 (J-009642-06-0002). The siRNAs at a concentration of 5 nM were reverse transfected using Lipofectamine RNAiMAX transfection reagent (Invitrogen, VIC, Australia) according to the manufacturer’s protocol. DECR1 downregulation (>80%) was confirmed on mRNA and protein levels.

### Generation of stable shDECR1 and hDECR1 LNCaP cells

*Short hairpin lentiviral expression vector*. LNCaP cells were transduced with the universal negative control shRNA lentiviral particles (shControl), DECR1 shRNA lentiviral particles (shDECR1) or hDECR1 (GFP-Puro) designed by GenTarget Inc. (San Diego, CA, USA) according to the manufacturer’s protocol. The sequence of the DECR1 shRNA and the hDECR1 is depicted in Figure S5.

### Functional Assays

#### Cell Viability

Cells were seeded in triplicates in 24-well plates at a density of 3.0 × 10^4^ – 6 × 10^4^ cells/well and reverse transfected with siRNA overnight. Cells were manually counted using a hemocytometer 96 hours post-siRNA knockdown and viability was assessed by Trypan Blue exclusion as described previously (81).

#### Cell Migration

Transwell migration assays were performed using 24-well polycarbonate Transwell® inserts (3422, Sigma Aldrich). LNCaP, 22RV1 and MR49F cells transfected overnight with siRNA were seeded into the upper chamber of the Transwell at a density of 8 × 10^4^ - 2.5 × 10^5^ cells/well in serum-free medium. 650 µl of medium containing 5% FBS was added to the bottom chamber. Cells were incubated at 37°C for 48 hours. Non-migrated cells were gently removed using a cotton-tipped swab. The inserts were fixed in 4% paraformaldehyde for 20 minutes and stained with 1% crystal violet for 30 minutes. The images of migrated cells were captured using the Axio Scope A1 Fluorescent Microscope (Zeiss) at 40X magnification. The number of migrated cells were counted manually and presented as percentages relative to control cells ± SEM.

#### Cell Invasion

Cell invasion were assessed using 24-well-plate BD Biocoat Matrigel Invasion Chambers (In Vitro Technologies, NSW, Australia) according to the supplier instructions. After 48 hours of siRNA transfection, 650 µl of medium containing 10% FBS was added to the bottom chamber, and equal number of cells within 1% FBS-contained medium were transferred to the upper chamber. After incubation at 37°C, 5% CO_2_ for 48 hours, non-invading cells as well as the Matrigel from the interior of the inserts were gently removed using a cotton-tipped swab. The inserts were fixed in 4% paraformaldehyde for 20 minutes and stained with 1% crystal violet for 30 minutes. The images of invaded cells were captured using the Axio Scope A1 Fluorescent Microscope (Zeiss) at 40X magnification. The number of invaded cells were counted manually and presented as percentages relative to control cells ± SEM.

#### Colony Formation Assay

DECR1 stable knockdown cells (shDECR1) or negative control cells (shControl) were prepared in a single-cell suspension before being plated in 6-well plates (500 cells/well). Cells were incubated for 2 weeks at 37°C and medium was replenished every 3-7 days. After 3 weeks, cells were washed with PBS, fixed with 4% paraformaldehyde and stained with 1% crystal violet for 30 minutes. Colonies were counted manually, and results were reported as number of colonies ± SEM.

#### 3D Spheroid Growth Assay

LNCaP and 22RV1 cells were transfected with siRNA in 6-well plates for 48 hours. Cells were collected and prepared at a concentration of 7.5 × 10^4^ cells/ml. Cell suspensions (1500 cells in 20 µl) were pipetted onto the inside of a petri dish lid, and 15ml of PBS was added to the dish to prevent the drops from drying. The petri dishes were reassembled and incubated at 37°C for 5 days. Photos of the formed spheres were captured, and the sphere volume was determined using ReViSP software (87).

### Seahorse Extracellular Flux Analysis

Cells were plated on XF96 well cell culture microplates (Agilent, VIC, Australia) at equal densities in substrate-limited medium (DMEM with 0.5mM glucose, 1.0mM glutamine, 0.5 mM carnitine and 1% FBS) and incubated overnight. One hour before the beginning of OCR measurement, the cells were changed into FAO Assay Medium (111 mM NaCl, 4.7 mM KCl, 2.0 mM MgSO_4_, 1.2 mM Na_2_HPO_4_, 2.5 mM glucose, 0.5 mM carnitine and 5 mM HEPES).

After baseline OCR is stabilized in FAO Assay Medium, 200 µM of linoleic-acid (LA) or palmitic acid (PA) were added before initializing measurements. Extracellular flux analysis was performed using the Seahorse XF Cell Mitochondrial Stress Test kit (Seahorse Bioscience) according to the manufacturer’s protocol. Extracellular flux experiments were performed on a Seahorse XF96 Analyzer and results were analysed using Seahorse Wave software for XF analyzers. The OCR values were normalized to cell numbers in each well.

### Metabolomics

LNCaP cells were transfected with siRNA for 96 hours in no glucose medium (containing 10% FBS) supplemented with 2.5 mM glucose in 6-well plates. Cells were washed with 1 ml of 37°C Milli-Q water on the shaker for 2 seconds. The plate was placed in sufficient volumes of liquid nitrogen, enough to cover the surface of the plate and was briefly stored on dry ice. 600 µl of ice cold 90% 9:1 methanol:chloroform (MeOH:CHCl_3_) extraction solvent containing the internal standards (0.5 µl/samples) was added onto each well and allowed to incubate for 10 minutes. Cells were collected into 1.5 ml Eppendorf tubes, incubated on ice for 5 minutes, and centrifuged at 4°C for 5 minutes at 16,100g. The supernatant was then transferred into a fresh 1.5 ml Eppendorf tube and allowed to dry in a Speedvac. Dried samples were derivatised with 20 µl methoxyamine (30 mg/ml in pyridine, Sigma Aldrich) and 20 µl N,O-Bis(trimethylsilyl)trifluoroacetamide (BSTFA) + 1% Trimethylchlorosilane (TMCS). The derivatised samples were analysed using GC QQQ targeted metabolomics as described in (88).

### Mitochondrial ROS Measurement

LNCaP cells were transfected with siRNA for 96 hours in 6-well plates. Cells were collected into fluorescence-activated cell sorting (FACS) tubes and stained with 2.5 mM of MitoSOX™ Red stain (Thermo Fisher Scientific, VIC, Australia) for 30 minutes in a 37°C water bath. Cells were centrifuged at 1,500rpm for 5 minutes, washed twice with 500 µl of PBS, and resuspended in 500 µl of pre-warmed PBS before the samples are read on a BD FACSymphony™ flow cytometer.

### Acyl-Carnitine Measurement

Total lipids were extracted from cells using two-phase extraction with methyl-tert-butyl-ether (MTBE)/methanol/water (10:3:2.5, v/v/v) (89). Cell pellets were frozen in 8:1 methanol/water prior to extraction with the above solvent mixture; for cell culture supernatant samples, the cell culture medium replaced the water component. Deuterated (D3)-palmitoylcarnitine was included as an internal standard (200 pmole/sample for LNCaP samples; 20 pmole/sample for tumor explant samples). Samples were reconstituted in 200 μL of the HPLC starting condition, defined below.

Acylcarnitines were quantified by liquid chromatography-tandem mass spectrometry using a Q-Exactive HF-X mass spectrometer with heated electrospray ionization and a Vantage HPLC system (ThermoFisher Scientific). Extracts were resolved on a 2.1 × 100 mm Waters Acquity C18 UPLC column (1.7 µm pore size), using an 18 min binary gradient at 0.28 mL/min flow rate, as follows: 0 min, 80:20 A/B; 3 min, 80:20 A/B; 6 min, 57:43 A/B; 8 min, 35:65 A/B; 9 min, 0:100 A/B; 14 min, 0:100 A/B; 14.5 min, 80:20 A/B; 18 min, 80:20 A/B. Solvent A: 10 mM ammonium formate, 0.1% formic acid in acetonitrile:water (60:40); Solvent B: 10 mM ammonium formate, 0.1% formic acid in isopropanol:acetonitrile (90:10). Data was acquired in positive ion mode with data-dependent acquisition (full scan resolution 70,000 FWHM, scan range 220–1600 *m/z*). The ten most abundant ions in each cycle were subjected to fragmentation (collision energy 30 eV, resolution 17,500 FWHM). An exclusion list of background ions was used based on a solvent blank. TraceFinder v5.0 (Thermo Fisher) was used for peak detection and integration, based on exact precursor ion mass (*m/z* tolerance 4 ppm) and *m/z* 85.0 acylcarnitine product ion. Peak areas were normalised to the D3-palmitoylcarnitine internal standard.

### *In Vivo* Experiment

LNCaP cells (5 × 10^6^ cells in 50μL 10% FBS/RPMI 1640 medium) were co-injected subcutaneously with 50μL Matrigel in 6-week-old Nod Scid Gamma male mice (Bioresource Facility, Austin Health, Heidelberg, Australia). When tumors reached ∼200mm^3^, mice were randomized in different therapy groups. One group was left untreated (*n* = 5), one group was treated with vehicle control (10% DMSO/PBS; *n* = 5), one group was treated with enzalutamide (10 mg/kg MDV3100 in 10% DMSO/PBS) and one group was castrated by surgical castration under isofluorane anaesthesia (*n* = 9). Five of the ten castrated mice were then treated daily with enzalutamide (10 mg/kg MDV3100 in 10%DMSO/PBS) by oral gavage for 7 days. Enzalutamide therapy of castrated mice started five days after surgery.

In a second study, DECR1 stable knockdown cells (shDECR1) or negative control cells (shControl) (5 × 10^6^ cells in 50μL 10% FBS/RPMI 1640 medium) were co-injected subcutaneously with 50μL matrigel in 6-week-old Nod Scid Gamma male mice. Tumours were measured using a calliper and their volumes were calculated using the formula length × width^2^/2.

For both studies, tumours were excised and half was snap frozen for RNA extraction while the other half was formalin fixed and paraffin embedded.

### Statistical Analysis

Results are reported as mean ± S.E.M. Statistical analysis was performed using GraphPad Prism (V7.0 for Windows). Differences between treatment groups were compared by T-test or one-way ANOVA followed by Tukey or Dunnett post hoc test. Significance is expressed as \**p* < 0.05, \*\**p* < 0.01, \*\*\**p* < 0.001, \*\*\*\**p* < 0.0001.

## Acknowledgments

The results published here are in part based on data generated by The Cancer Genome Atlas, established by the National Cancer Institute and the National Human Genome Research Institute, and we are grateful to the specimen donors and relevant research groups associated with this project. Tissues for the patient-derived explants used in the study were collected with informed consent via the Australian Prostate Cancer BioResource and we thank the doctors, patients and health care professionals involved. We acknowledge expert technical assistance in the study from Natalie Ryan, Joanna Gillis, Kayla Bremert, Samira Khabbazi, Nhi Huynh, and Holly P. McEwen. Flow cytometry analysis was performed at the South Australian Health Medical Research Institute (SAHMRI) in the ACRF Cellular Imaging and Cytometry Core Facility. The Facility is generously supported by the Australian Cancer Research Foundation (ACRF), Detmold Hoopman Group and Australian Government through the Zero Childhood Cancer Program. The authors thank Metabolomics Australia, Bio21 Institute, and Adelaide Microscopy (University of Adelaide).

## Author Contributions

Conception and design: ZDN, JVS, LMB. Development of methodology: ZDN, CYM, JD, IB, MMC, ASD, DJL, LAS. Acquisition of data: ZDN, CYM, JD, IB, SI, ASD, RKS, MM. Analysis and interpretation of data: ZDN, CYM, IB, SI, MM, ASD, LGH, DJL, LAS, AJH. Writing the manuscript: ZDN, CYM, LMB Review and editing of the manuscript: All authors

