## Supplementary Figures and Tables for "DECR1 is an androgen-repressed survival factor that regulates PUFA oxidation to protect prostate tumor cells from ferroptosis"

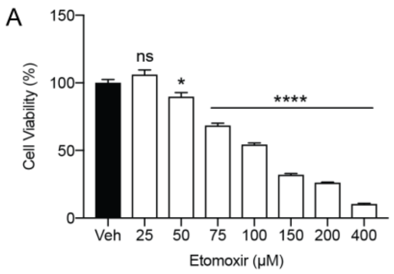


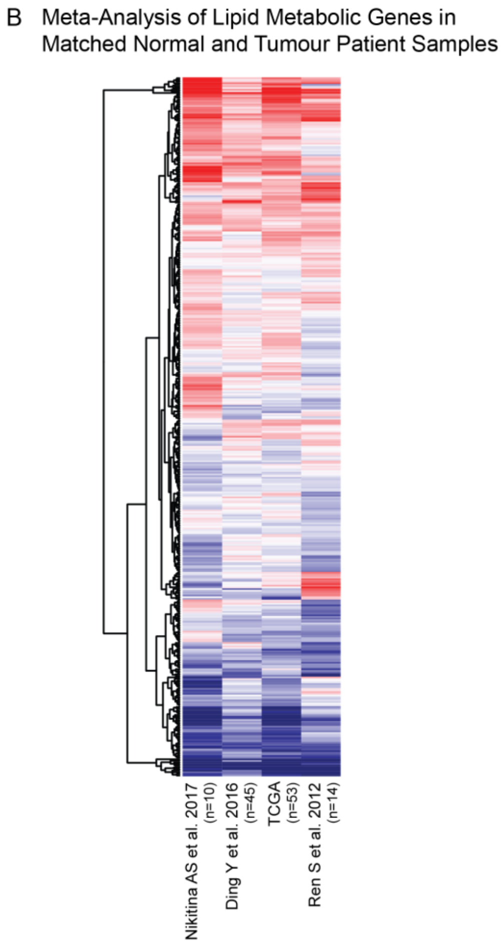


C

| Dataset | Fold Change | P value | No. of Tissues | |
|---|---|---|---|---|
| | | | Normal | Malignant |
| TCGA | 1.44 | 0.0001 | 499 | 51 |
| Ding | 1.12 | 0.0002 | 45 | 45 |
| Tomlins | 2.389 | 1.81E-04 | 23 | 30 |
| Luo | 1.404 | 0.001 | 15 | 15 |
| Vanaja | 1.9 | 5.57E-07 | 15 | 15 |
| Grasso | 1.311 | 5.25E-05 | 28 | 59 |
| Areedouani | 1.311 | 9.00E-03 | 8 | 13 |
| Lapointe | 1.068 | 0.062 | 41 | 62 |
| Singh | 2.294 | 2.55E-04 | 50 | 52 |
| Weslh | 1.419 | 1.00E-03 | 9 | 25 |

**Figure.S1. (A)** Cell viability of LNCaP cells treated with indicated concentrations of etomoxir for 72 hours. Cell viability were determined using CyQUANT® Cell Assay**. (B)** A meta-analysis of 735 lipid metabolism genes using four clinical datasets with malignant and matched normal RNA-sequencing data (n=122). Genes were rank-ordered on the basis of their *meta effect size* scores in PCa malignant tissues versus matched normal. **(C)** Fold change of DECR1 mRNA expression in malignant tissues compared to benign/normal tissues. Data were obtained from Oncomine.

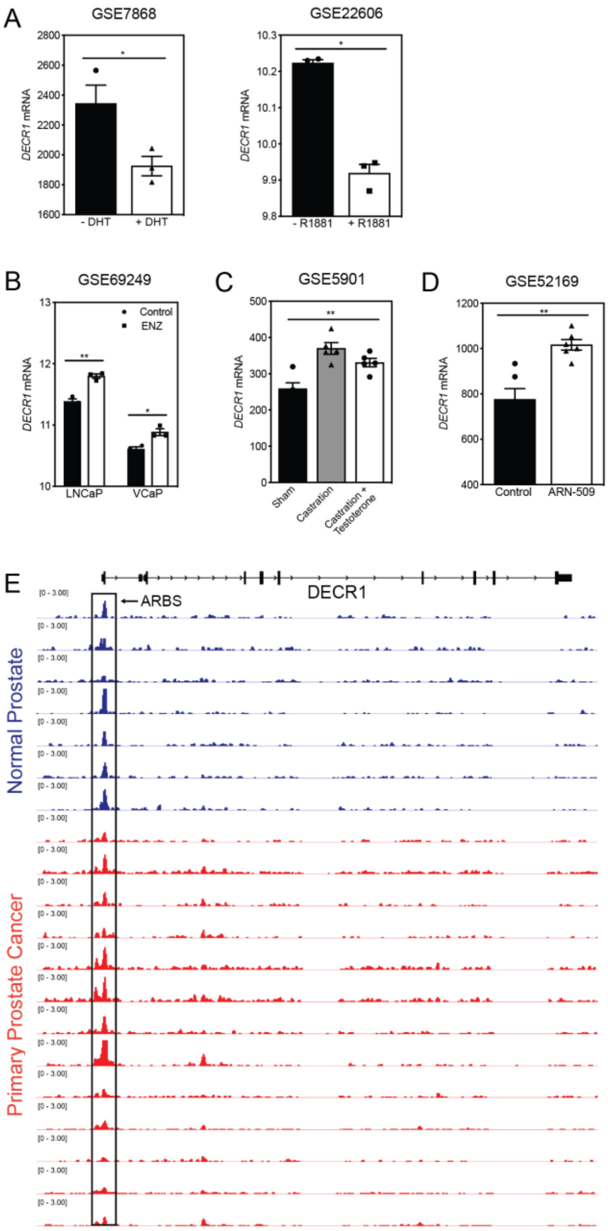


**Figure.S2. (A)** Bar graphs of DECR1 mRNA expression in two publically available datasets shows DECR1 mRNA expression decreased after LNCaP treatment with DHT (GSE7868) or R1881 (GSE22606). **(B)** DECR1 expression increased in LNCaP and VCaP upon treatment with Enzalutamide (GSE69249). **(C)** DECR1 mRNA expression increased in mouse prostate gland after castration but decreased after testosterone administration. **(D)** ARN-509 (Apalutamide) treatment of LNCaP/AR xenograft increased DECR1 mRNA expression (GSE52169). **(E)** AR ChIP-sequencing data from normal human prostate and primary human prostate tumour specimens (normal, n=7; tumour, n=13). Data from GSE56288. Statistical analysis was performed using two-tailed Student's *t*-test: **p*<0.05, ***p* <0.01 and *****p* <0.0001.

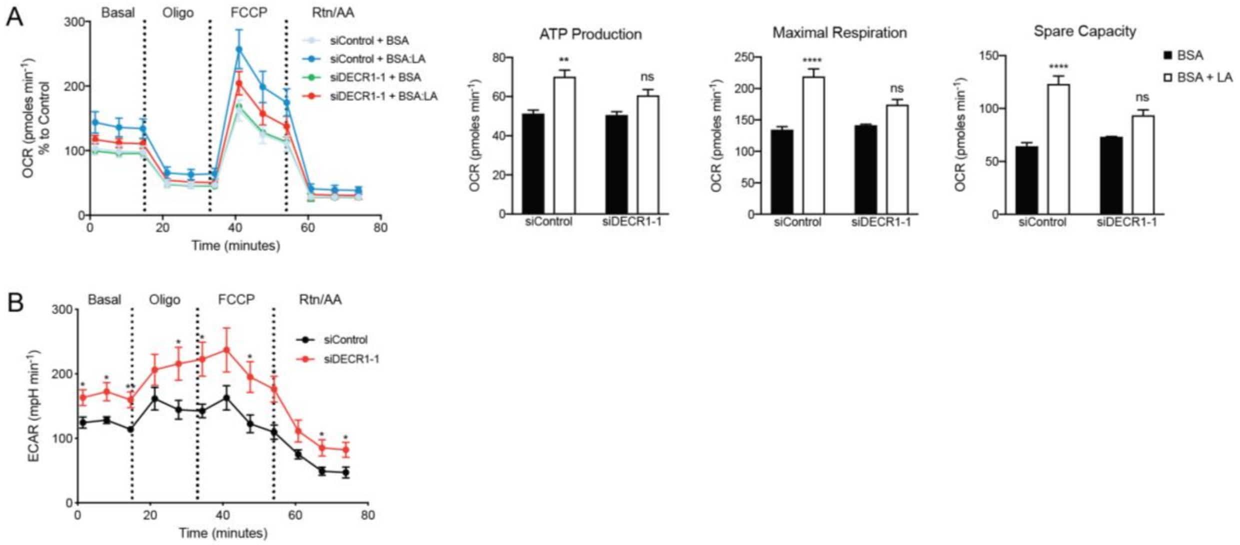


**Figure.S3. (A)** Oxygen consumption rate (OCR) was assessed in LNCaP cells supplemented with the PUFA linoleic acid (LA). Each data point represents an OCR measurement. ATP production, maximal mitochondrial respiration and mitochondrial spare capacity were assessed. **(B)** Extracellular acidification rate (ECAR) was assessd in LNCaP cells. Each data point represents an ECAR measurement. Statistical analysis was performed using two-tailed Student's *t*-test: **p*<0.05, ***p* <0.01 and *****p* <0.0001.

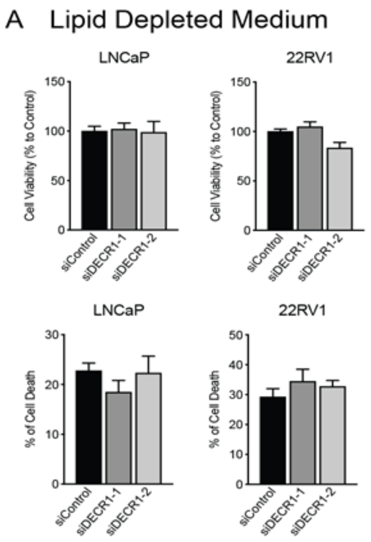


**Figure.S4. (A)** Cell viability and cell death after DECR1 knockdown in LNCaP and 22RV1 cultured in full serum media. Cell viability and cell death were measured using trypan blue exclusion following 96 hours DECR1 knockdown. Percentages are represented relative to the control siRNA; *n* = 3 independent experiments per cell line.

**shControl sequence:** GTCTCCACGCGCAGTACATTT
cloned shRNA hairpin sequence:

GTCTCCACGCGCAGTACATTTcgagAAATGTACTGCGCGTGGAGAC

**shDECR1-3 sequence:** GTGATTCAACCAGGGCCTATA
cloned shRNA hairpin sequence:

GTGATTCAACCAGGGCCTATAcgagTATAGGCCCTGGTTGAATCAC

blue: sense
black: loop
red: anti-sense

**hDECR1**
sub-cloned human target coding sequences:

Human DECR1
atgtcaggactggggaagaagcatctcctgctcatgggggagtttttcagttatgggacaaaaatattatatcaaaacactgaagcttt
gcaatctaaattcttttcacctcttcaaaaagcgatgctaccacctaatagttttcaaggaaaagtggcattcattactgggggaggtact
ggccttggtaaaggaatgacaactcttctgtccagcctaggtgctcagtgcgtgatagccagccggaagatggatgttttgaaagctac
cgcagaacaaatttcttctcaaactggaaataaggttcatgcaattcagtgtgatgtgagggatcctgatatggttcaaaacactgtgtc
agaactgatcaaagttgcaggacatcctaatattgtgataaacaatgcagcagggaattttatttctcctactgaaagactttctcctaat
gcttggaaaaccataactgacatagttctaaatggcacagccttcgtgacactagaaattggaaaacaactaattaaagcacagaaag
gagcagcatttctttctattactactatctatgctgagactggttcaggttttgtagtaccaagtgcttctgccaaagcaggtgtggaagcc
atgagcaagtctcttgcagctgaatggggtaaatatggaatgcgattcaatgtgattcaaccagggcctataaaaaccaaaggtgcctt
tagccgtctggacccaactggaacatttgagaaagaaatgattggcagaattccctgtggtcgcctggggactgtagaagaactcgca
aatcttgctgctttcctttgtagtgattatgcttcttggattaatggagcagtcattaaatttgacggtggagaggaagtacttatttcaggg
gaattcaacgacctgagaaaggtcaccaaggagcagtgggacaccatagaagaactcatcaggaagacaaaaggttcc

**Figure.S5. The sequence of the DECR1 shRNA and the hDECR1**

**Supplementary Table 1: Primary antibody and stains details**

| Antibody/stain | Catalogue number | Producer/Supplier |
|---|---|---|
| **β-Actin** | A5441 | Sigma-Aldrich (NSW, Australia) |
| **HSP90** | 48745 | Cell Signalling Technology (Genesearch, QLD, Australia) |
| **DECR1** | HPA023238 | Prestige Antibodies (Sigma-Aldrich, NSW, Australia) |
| **Malondialdehyde** | ab6463 | Abcam (VIC, Australia) |
| **Androgen receptor** | sc-816 | Santa Cruz Biotechnology (MetaGene, QLD, Australia) |
| **MitoTracker Red CMXRos** | M7512 | Thermo Fisher Scientific (VIC, Australia) |
| **MitoSOX™ Red Mitochondrial Superoxide Indicator** | M36008 | Thermo Fisher Scientific (VIC, Australia) |
| **KI67** | M7240 | DAKO (Agilent Technologies Australia, VIC, Australia) |

**Supplementary Table 2: Primer sequences used in qPCR**

| | | |
|---|---|---|
| **DECR1** | ***sense*** | CTAAATGGCACAGCCTTCGT |
| | ***Antisense*** | AACCTGAACCAGTCTCAGCA |
| **GAPDH** | ***sense*** | TGCACCACCAACTGCTTAGC |
| | ***Antisense*** | GGCATGGACTGTGGTCATGAG |
| **PPIA** | ***Sense*** | GCATACGGGTCCTGGCAT |
| | ***Antisense*** | ACATGCTTGCCATCCAACC |
| **TUBA1B** | ***sense*** | CCTTCGCCTCCTAATCCCTA |
| | ***Antisense*** | CCGTGTTCCAGGCAGTAGA |
| **MKI67** | ***sense*** | GCCTGCTCGACCCTACAGA |
| | ***Antisense*** | GCTTGTCAACTGCGGTTGC |
| **L19** | ***Sense*** | TGCCAGTGGAAAAATCAGCCA |
| | ***Antisense*** | CAAAGCAAATCTCGACACCTTG |
| **GUSB** | ***Sense*** | CGTCCCACCTAGAATCTGCT |
| | ***Antisense*** | TTGCTCACAAAGGTCACAGG |

**Supplementary Table 3: Primer sequences used in ChIP-qPCR**

| | | |
|---|---|---|
| **DECR1** | ***sense*** | TTCTGGAGCGCTAAGAGAGC |
| | ***Antisense*** | AGGGCTTCATCTGACAGTGG |
| **KLK3** | ***sense*** | GCCTGGATCTGAGAGAGATATCATC |
| | ***Antisense*** | ACACCTTTTTTTTTCTGGATTGTTG |
| **NC2** | **sense** | GTGAGTGCCCAGTTAGAGCATCTA |
| | **Antisense** | GGAACCAGTGGGTCTTGAAGTG |
